# Mapping of Multilineage Tumor Cell Populations in Mouse Bladder Cancer

**DOI:** 10.1101/2024.12.04.626806

**Authors:** Nadine Schrode, Yang Hu, Haocheng Yu, Matthew Galsky, John Sfakianos, Robert Sebra, Kristin G. Beaumont, David J Mulholland

## Abstract

In bladder cancer (BLCA), the cellular expression of lineage markers with predictive biomarker function can have important implications for tumor progression, treatment response, and survival. However, tumor heterogeneity and the coexistence of distinct tumor subpopulations can complicate the utility of using a primary, clinically assigned tumor signature. In this report, we have applied the reference atlas, *Tabula Sapiens,* and deep learning model, UniCell, to conduct in-depth transcriptional analysis of unique lineage marker-defined cell clusters in a carcinogen (BBN) induced bladder cancer model. UniCell deconvolution has identified tumor populations, including urothelial, adenocarcinoma, squamous cell carcinoma, and mesenchymal tumor populations, each with cell-intrinsic gene expression signatures relevant to human BLCA progression. The identified tumor clusters contain uniquely basal, luminal, stromal, or hybrid cells in cell lineage marker expression. To understand the significance of these populations during progression, we used trajectory and pseudo-time analysis to show that cells uniquely basal are plastic in lineage identity and can evolve to form tumor populations composed of other lineage marker-defined signatures. Finally, pathway and drug enrichment analysis of tumor cell clusters were used to identify therapeutics that may preferentially target the identified tumor cell populations. *These data collectively define a molecular template that may uniquely profile molecular plasticity occurring during progression and response to therapy with important implications for human disease*.

**Significance:** Tumor heterogeneity is a mechanism for treatment resistance. Our study defines unique tumor subpopulations having differential therapeutic sensitivities and potential for lineage plasticity. Our modeling may impact the treatment of BLCA patients.

## INTRODUCTION

Lineage marker expression in tumor cells can provide predictive information about progression, response to therapy, and survival outcome [1] [2]. In human bladder cancer, basal and luminal phenotypes have been used to establish a quantitative algorithm and immunohistochemistry classifiers leading to the identification of clinical classification of basal-positive, luminal-positive, and hybrid-positive tumor cells and those that are double negative for both markers [3]. Double-negative tumor populations also include a portion of cells in an activated EMT state with an upregulation of transcription factors driving EMT and a downregulation of homotypic adhesion proteins [3]. Indeed, in human bladder cancer, EMT-like cells are associated with poor prognosis, while an umbrella subtype is related to positive survival outcomes [4].

Since the lineage composition of tumor cells is essential for therapeutic response, changes in lineage identity through plasticity can have important clinical implications [5-7]. Recent studies showing retinoid-induced basal to luminal plasticity and an associated tumor progression in mice [8] exemplify the capacity for lineage plasticity.

However, modeling and functional understanding of how different cell populations respond to treatment are complex. They require tumor models with well-defined molecular landscapes and the capacity to relate therapies to changes in these landscapes. Preclinical bladder cancer models initiated from carcinogens found in tobacco smoke produce mutationally driven primary tumors with progression and pathological kinetics like human disease [9, 10].

Our previous studies studied transcriptional heterogeneity and demonstrated the coexpression of multi-lineage markers in mouse and human tumor cells [11]. This report applies *the Tabula Sapiens* transcriptional atlas and UniCell deep learning to establish a detailed molecular landscape of primary BBN tumors, focusing on lineage marker expression with biomarker importance. We further apply drug prediction algorithms to identify drugs that may be effective alone or in combination with standard-of-care therapeutics for targeting signature pathways found.

## METHODS

### Ethics approval and consent to participate

Tumor specimens utilized to generate single-cell RNA sequencing data were derived from patients participating in project #10-1180 approved by the Icahn School of Medicine Institutional Review Board. Written informed consent was obtained from participants. The research conformed to the Declaration of Helsinki.

### Animal Experiments

All studies were conducted under the approved IACUC protocol LA13-00060.

### Sex as a biological variable

Our study data sets are derived from male mice with bladder tumors.

### Key Resources

#### Generation of carcinogen-induced mouse bladder tumors

We generated primary muscle-invasive bladder cancers (MIBCs) in adult FVB/NJ mice (n = 3) by administering the BBN carcinogen water (0.1%) for 14 weeks, followed by regular drinking water for four weeks. Tumor production and the resulting single-cell RNA seq data sets were processed according to previously conducted protocols [11] **(Fig. 1A)**.

**Fig. 1.**
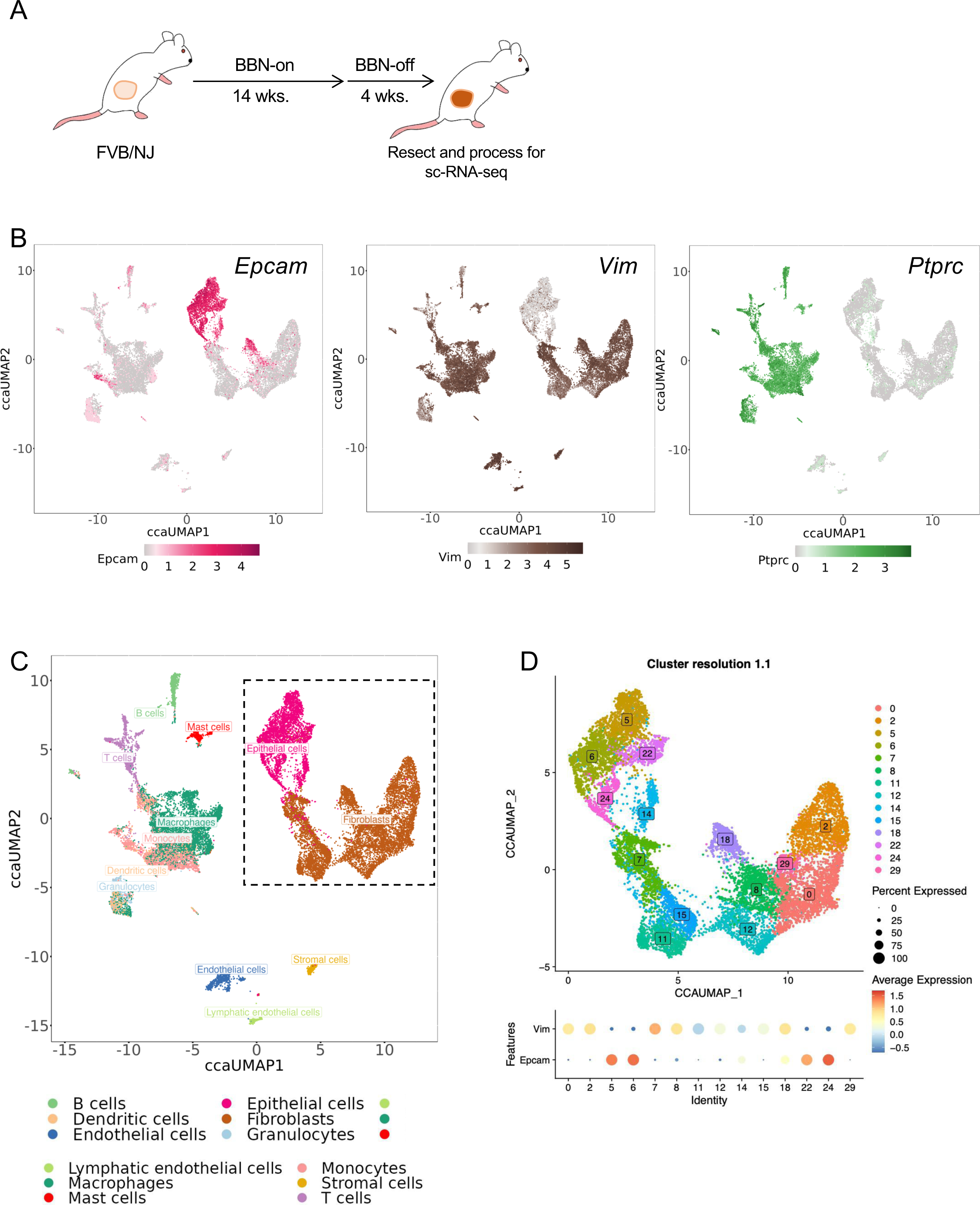
Single-cell transcriptomic profiling of BBN-induced mouse bladder tumors. (A) The BBN carcinogen is administered to FVB/NJ CAGS-luciferase mice to induce primary bladder tumors. (B) ccaUMAP plots showing markers for epithelial (Epcam, *Epcam*, red), stromal (Vimentin, *Vim*, brown), and immune cells (CD45, *Ptprc*, green). (C) Tumor cell populations are shown as integrated UMAP plots (ccaUMAP) derived from aggregate data sets (n=3) of BBN-induced tumors. Cell types are referenced according to the color-coded key. (D) Unsupervised clustering of tumor cell populations showing 14 unique clusters (integrated_snn_res.1.1) from the gated dashed box in C. Bubble plots show the expression of Vimentin and Epcam in each cluster. Relates to **Tables 1-2**.

### Single-cell transcriptome analysis

Single-cell RNA-seq data was pre-processed with scater [12] and normalized by scran [13]. Data integration, unsupervised cell clustering, and differential expression analysis were conducted using Seurat v3.0 [14]. Cells with more than three median absolute deviations were removed as outliers. Cells with less than 400 genes or, 1000 UMIs or more than 15% of mitochondria genes were filtered out from the analysis. The filtered data contained 27,998 cells and 24,421 genes from 6 samples. Cell-specific biases were normalized with pool-based size factors. The top 3000 highly variable genes were selected using the expression and dispersion (variance/mean) of genes, followed by a canonical correlation analysis (CCA) to identify common sources of variation between the patient and normal datasets. The first 105 CCA results were chosen to generate dimensional t-distributed Stochastic Neighbor Embedding (tSNE) plots, Uniform Manifold Approximation and Projection (UMAP) plots, and cell clustering by a shared nearest neighbor (SNN) modularity optimization-based clustering algorithm.

Cell types were manually identified by marker genes [15] and confirmed by reference-based cell type annotation generated by the SingleR (Single-cell Recognition) package using 358 mouse RNA-seq [16].

UniCell (10) was used to carry out a more detailed annotation of an epithelial-stromal subset. Differential expression analysis was performed using the MAST (Model-based Analysis of Single Cell Transcriptomics) algorithm [17].

Gene expression signatures were added by calculating the average expression levels of a set of genes, subtracted by the aggregated expression of control gene sets. All analyzed genes were binned based on averaged expression, and 100 control genes were randomly selected from each bin.

Trajectories were calculated using the Monocle3 [18] ‘learn graph()’ function and respective pseudo time was inferred using the population with the highest density of basal marker gene-expressing cells (Cluster 6). Genes differentially expressed across individual branches of the single-cell trajectory were identified through Moran’s I test (a measure of multi-directional and multi-dimensional spatial autocorrelation).

For over-representation (ORA) enrichment analysis in drug targets, the Drug Signature Database gene sets were used (https://dsigdb.tanlab.org/DSigDBv1.0/collection.html), with mechanisms of action sourced from the Broad’s drug repurposing hub (https://repo-hub.broadinstitute.org/repurposing-app). Pathway enrichment analysis used Reactome Pathways gene sets (https://reactome.org/download-data).

### Code availability

Scripts used for analysis are available at https://github.com/nyuhuyang/scRNAseq-BladderCancer.

## RESULTS

We previously studied lineage heterogeneity in mouse and human bladder tumors using single cell transcriptomics and immunohistochemistry [11]. Our current study has used the *Tabula Sapiens* atlas and UniCell deep learning to extend these observations and conduct a detailed analysis of the molecular landscape in the BBN mouse model of bladder cancer. We provide evidence for progression-dependent plasticity as well as differential pathway and drug dependencies as a function of unique subpopulation signatures. The following data support these conclusions.

### *Tabula Sapiens* and UniCell identify unique tumor clusters

Using unsupervised clustering of tumor cell populations, we used *Tabula Sapiens* [19], a human cell atlas of 500,000 cells from 24 organs, to define the transcriptional landscape of mouse cells isolated from muscle-invasive primary bladder tumors induced by the BBN carcinogen. Tumor cell types were classified using *Tabula Sapiens* and UniCell with aggregate numbers shown by cell type number **Table 1** (relates to Fig. 1) and cell cluster **Table 2** (refers to Fig. 1).

### Tumor populations hybrid in lineage marker expression are prevalent in BBN tumors

We dosed FVB/NJ mice with the BBN carcinogen and isolated tumor cells for single cell RNA-seq processing **(Fig. 1A)**. We assessed aggregate single-cell data sets (tumors 4950, 8524, 8525) to further understand the molecular heterogeneity of untreated primary tumors including the presence of tumor cells hybrid in expression for lineage biomarkers. For this, we evaluated the expression of a pan epithelial (EPCAM-*Epcam*, red), a pan stromal marker (VIM-*Vim*, brown), and a widely expressed immune cell marker (CD45-*Ptprc*, green) **(Fig. 1B)**. Both Epcam and Vim were broadly expressed including multiple clusters having high coexpression in epithelial and fibroblastic like cells. Conversely, CD45-*Ptprc* gene expression was restricted to immune cells **(Fig. 1B-C)**. From here, we focused exclusively on tumor cells with epithelial and fibroblastic marker expression (dashed gated populations) using high-resolution UMAP plots (integrated_snn_res.1.1) to identify 14 transcriptionally unique cell clusters **(Fig. 1C-D, dashed gate)**. The identified clusters were color-coded and ranked by cell numbers, where cluster 0 had the highest number of cells and cluster 29 had the lowest numbers shown as aggregate data.

We generated bubble plots for each identified cluster to relate clusters with *Epcam* and *Vim* expression. We observed clusters 5, 6, 22, and 24 to be *Epcam*-high and clusters 0, 2, 7, 8, 12, 15, 18 and 29 to be *Vim*-high. Clusters 14 and 18 showed high coordinate expression of *Epcam* and *Vim* **(Fig. 1D), Table 2** (relates to Fig. 1).

*Together, these data define 14 unique tumor subpopulations, including those with hybrid epithelial and mesenchymal lineage marker expression*.

### UniCell deep learning classification model for cell type prediction

Next, we used UniCell deep learning to evaluate cell type signatures related to each of the 14 unique clusters identified. Unicell is a deep learning model pre-trained on the world’s largest fully integrated scRNA-seq training database, comprising over 28 million single cells spanning more than 840 cell types from 899 studies [20]. UniCell predicts the signature probability of each cell cluster and best matches by computing attribution scores for all input genes. It associates the expression of each gene with the cell type prediction with classification genes indicated for each cell type in **Table 3** (relates to Fig. 2). Unicell identified cellular signatures with urothelial qualities (clusters 5, 6, 14, 22, 24), adenocarcinoma (cluster 7), squamous carcinoma (clusters 11, 15), fibroblasts (clusters 0, 8, 12, 29), mesenchymal stem cells (cluster 2) and epicardial cells (cluster 18) **(Fig. 2A)**.

**Fig. 2.**
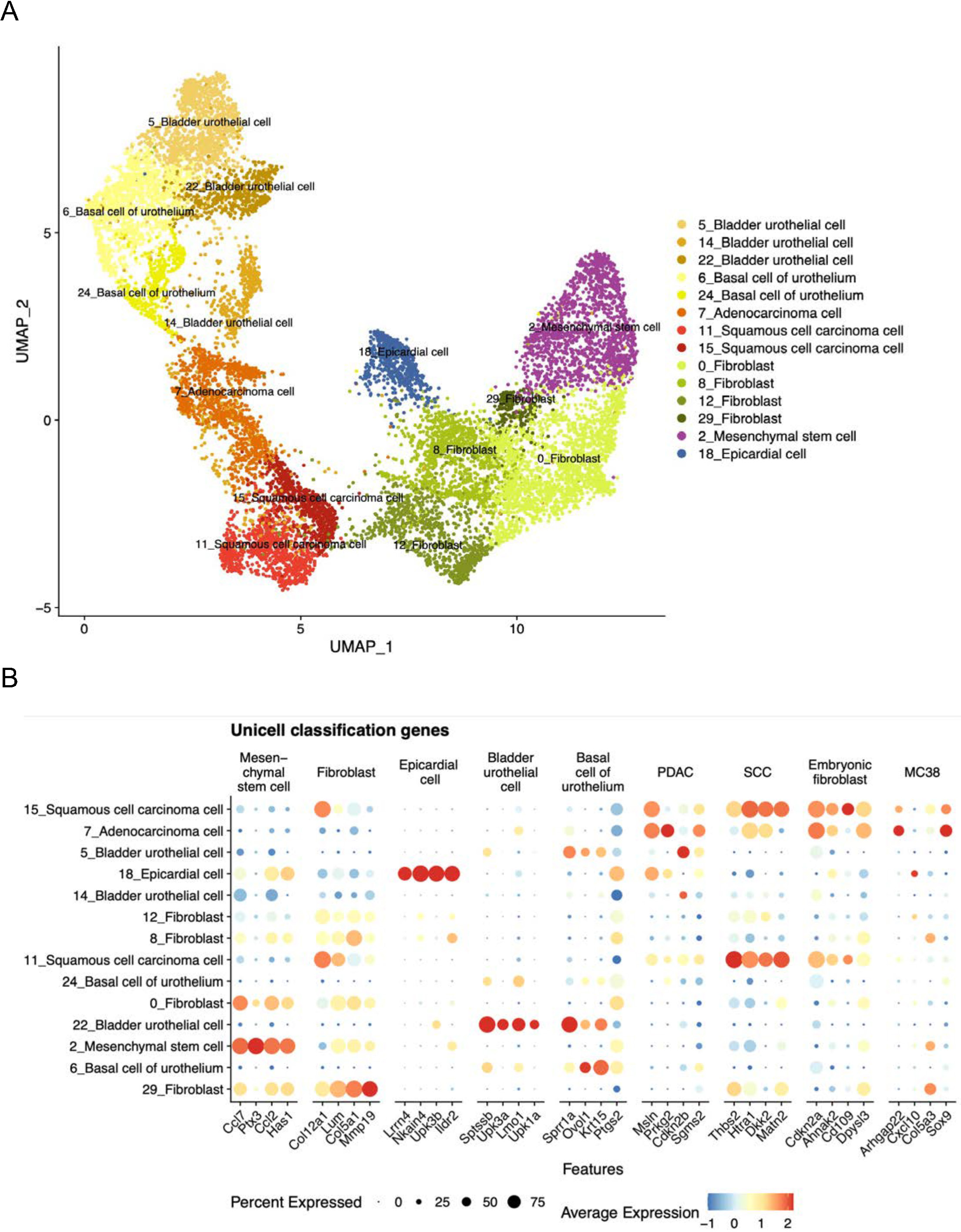
Bladder tumor cell types shown by UniCell classification. (A) UMAPs show 14 cell clusters classified into nine cell type signatures by UniCell gene classification. (B) Bubble plots showing the expression of the top 4 classification genes for each cell type. Relates to **Tables 3-6**.

We identified nine signatures represented by the top four classification genes to define unique expression patterns in each cluster **(Fig. 2A)**. A complete ranking of Unicell genes for each signature is shown in **Table 4** (relates to Fig. 2). Classification signatures were found in all clusters as unique or multiple entities. For example, mesenchymal stem cells were defined predominantly by a single Unicell signature (Ccl2, Ptx3, Ccl2, Has1), but others, such as squamous cell carcinoma (SCC) cells, showed high expression of Unicell classifiers belonging to PDAC, SCC, embryonic fibroblast and MC38 **signatures (Fig. 2B)**. Notably, Unicell analysis showed that BBN primary tumors contain subpopulation signatures representative of most of the major variants of human bladder cancer, including urothelial carcinoma (clusters 5, 6, 14, 22, 24), squamous cell carcinoma (cluster 11, 15), adenocarcinoma (cluster 7), and potentially small cell carcinoma.

Differential Pathway enrichment analysis was conducted using gene IDs shown in column G of **Table 5** (relates to Fig. 2). In “Bladder Urothelial Cells,” pathways with significant enrichment included Zinc transporters, XBP1 chaperones, YAP signaling, and WNT5A dependent signaling. Differently, “Basal Cells of Urothelium” included pathways related to adhesion, keratinization, and markers of homologous recombination. The “adenocarcinoma” clusters showed pathway enrichment of genes associated with cell cycle checkpoints and DNA synthesis while “squamous cell carcinoma” (SCC) was most enriched for pathways related to abnormal metabolism and glycosylation.

To consider whether differential gene expression signatures from individual cell types could regulate therapeutic response, we conducted an over-representation enrichment analysis (ORA). Therapeutics identified which may preferentially target bladder (1) bladder urothelial cells: pentoxifylline and dipyridamole – a drug known to sensitize the potential of immune and chemotherapies; (2) bladder adenocarcinoma: carfilzomib – a proteosome inhibitor and (3) bladder squamous cell carcinoma: Sorafenib – a multi kinase inhibitor, and dasatinib – a SRC kinase inhibitor. **Table 6** (relates to Fig. 2)

*These data define unique tumor subpopulations in the BBN tumor model relevant to major pathological states of human bladder cancer. Enrichment analysis identifies differentially activated pathways in each major subcluster and potential sensitivities to drugs*.

### Prevalence of multilineage marker positive cells in BBN primary tumors

We used Unicell to map clusters containing cells with coordinate expression of basal, luminal, and mesenchymal lineage biomarkers. Classifier genes that comprise basal **(Fig. 3A, Fig. S2)**, luminal **(Fig. 3B, Fig. S3),** and EMT signatures **(Fig. 3C, Fig. S4)** are shown as UMAP plots (left), bubble plots of total gene signatures (middle), and bubble plots of individual genes (right). We observed heterogeneity in gene expression for each signature, in the percent of cells per cluster expressing each gene and in the average gene expression per cluster. For example, for basal genes, Krt14, Krt5, Krt6a, and Trp63 were expressed highly in clusters 5, 6, and 24 compared to Krt1 and Krt6b **(Fig. 3A).**

**Fig. 3.**
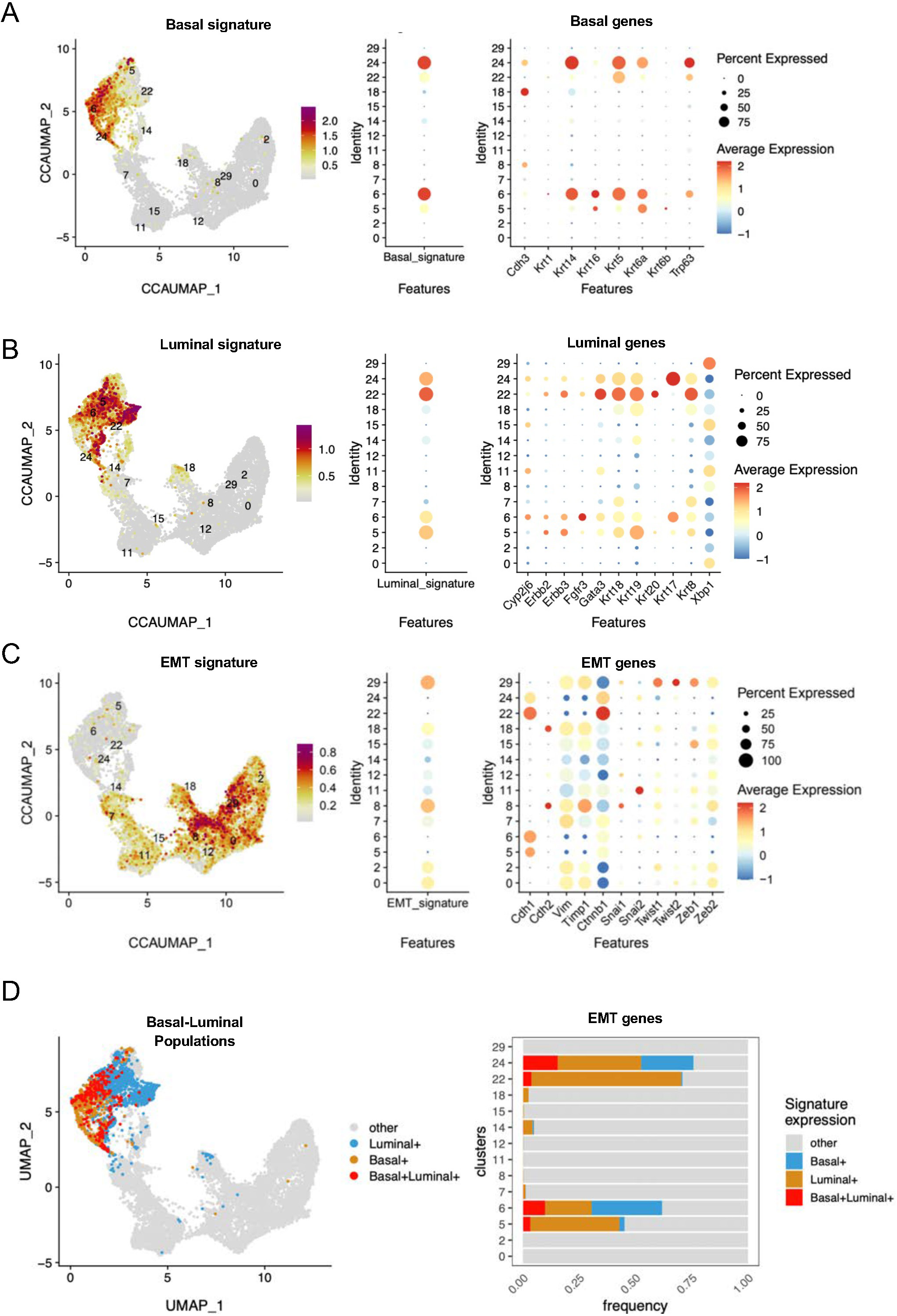
Coordinate cell expression of basal, luminal, and EMT lineage marker-positive cells. UMAP and bubble plots show signatures for (A) basal, (B) luminal, and (C) stromal markers. (D) Annotation of basal, luminal, and basal-luminal marker positive populations based on their gene signature expression (left) and expression of EMT markers in these compartments (right).

Using these signatures, we identified clusters 5, 6, 22, and 24 as having a heightened basal signature defined by Cdh3, Krt14, Krt16, Krt5, Krt6a, and Trp63 expression **(Fig. 3A**). High luminal gene expression was detected in clusters 5, 6, 7, 18, 22, 24 and 29 **(Fig. 3B).** EMT signatures occur in many solid tumors, including human bladder cancer [21]. Thus, we assessed basal or luminal marker-positive tumor cells for concomitant expression of EMT signature genes. We found clusters 0, 2, 8, 18, and 29 **(Fig. 3C, left)** to have a high percentage of cells with high average EMT marker expression **(Fig. 3C, middle)**. EMT genes, including Vim, Timp1, Ctnnb1, Twist1, Zeb1, and Zeb2, presented high average expression in many clusters **(Fig. 3C, right)**. While we detected less overlap between basal and EMT markers, we identified cluster 7 to have heightened co-expression of luminal (Krt8, Krt18) and EMT (Vim, Timp1, Ctbbn1, Twist1, Twist2, Zeb1, Zeb2) expression. Color coded UMAP plots and histogram analysis showed that basal-luminal double-positive cells resided in clusters 5, 6, 18, 22, and 24 **(Fig. 3D, Fig. S1)**. Of interest is cluster 18, which showed basal (Cdh3, Krt15), luminal (Krt18, Krt19, Krt8, Xbp1) and EMT (Vim, Timp1, Ctnnb1, Twist, Zeb2) coexpression. E-Cadherin (Cdh1), which is often lost during EMT, was found in reduced expression in multiple clusters and associated with heightened expression of Beta-catenin (Ctnnb1), a closely related protein critical adhesion both in urothelial clusters (5, 6, 22, 24) but also in adenocarcinoma (cluster 7) and other squamous cell and fibroblast clusters. Widespread expression of EMT transcription factors known to promote EMT reprogramming, including Snail1, Snail2, Twist1, Twist2, Zeb1, and Zeb2, was also identified.

*This data supports the existence of unique cellular clusters characterized by a steady-state bi-lineage and potentially multi-lineage marker expression*.

### *‘*Trajectory analysis predicts lineage plasticity of tumor subpopulations

Our study [11] and previous investigations [22] showed that Krt5-positive cells can function as a cellular origin for carcinogen-induced tumorigenesis. Thus, we hypothesized that basal cell clusters identified in this study (positions C6, C24) could form other clusters, including luminal and mesenchymal signatures, when subject to pseudotime analysis.

We generated trajectory graphs initiated from node 1 (white circle) **(Fig. 4A)** and plotted the highest density of basal cells initiated from this cellular location as a function of trajectory plots **(Fig. 4B).** Using pseudo-time analysis, we evaluated gene expression patterns that began from node 1 that terminated at different lineage marker defined clusters (C5, C22, C11, C2, C8/0). With these different gene expression states, we assessed the change in lineage marker expression over time **(Fig. 4C)**, including trajectories from basal signature genes C6 to C5 **(Fig. 4D)**, luminal signature genes C6 to C22 **(Fig. 4E)**, luminal genes from C6 to C11 (squamous cell carcinoma) **(Fig. 4F)**, and EMT genes from C6 to C11 (squamous cell carcinoma) **(Fig. 4G)**.Initially, the expression changes of known basal, luminal, and EMT biomarkers were assessed along trajectory branches **(Fig. 4D-G)**. For example, in C6 (basal) to C5 (luminal), we measured marked reductions in Trp63 and Chd3. During the trajectory progression from C6 to luminal C22 cluster, Krt20, Krt18, and Gata3 increased while Krt17 decreased. From C6 to C11 (SCC), Fgfr3, Krt19, Krt17 and Krt8 were reduced while Vim, Snail1, Snail2 and Zeb1 were increased **(Fig. 4H)**.

**Fig. 4.**
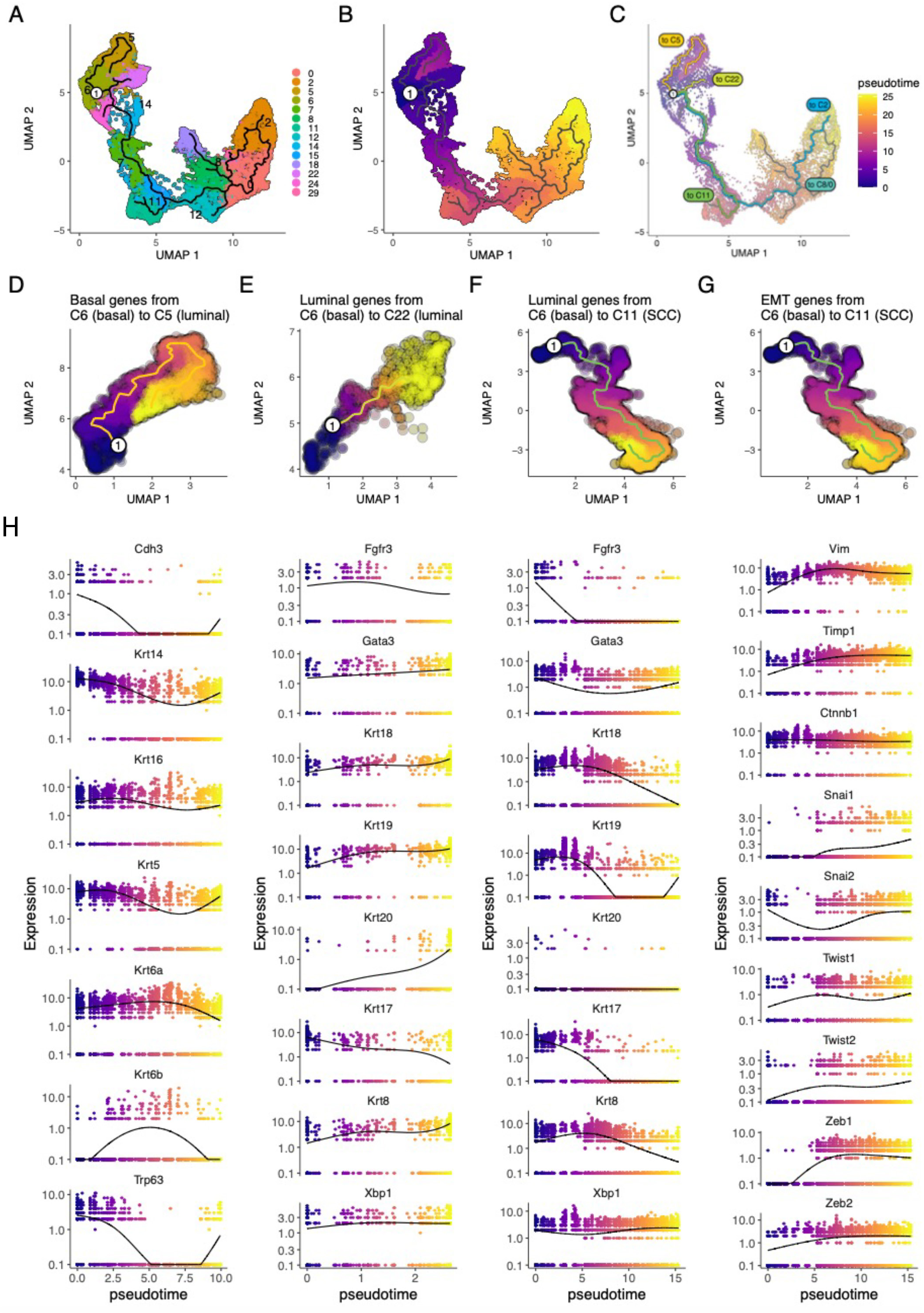
Trajectories and differentially expressed genes (DEGs) during pseudotime analysis initiated from basal cells. (A) Trajectory graphs plotted for 14 cell clusters. (B) The pseudotime plot, with the highest density of basal cells (node 1, white circle), indicating the starting point for trajectory analysis. (C) Trajectory graphs and pseudotime plots, superimposed with color-coded trajectory segments, show the start (C6) and termination points defined by unique clusters (C5, C22, C11, C22, C8/0). Expression of DEGs along pseudotime for (D) basal, (E-F) luminal, and (G) EMT gene in relevant clusters. Relates to **Table 7**.

*The analysis supports statistical differences (p values, Moran’s Test) between genes of interest along pseudotime trajectory branches, as shown in* ***Table 7*** *(relates to Fig. 4)*.

### Differentially Expressed Genes (DEGs) during pseudotime along segments

Pseudotime analysis supports the presence of transcriptional changes between tumor clusters during progression and lineage differentiation. To identify potential unknown targets relevant to progression, lineage plasticity, and therapeutic response we assessed DEGs of unknown targets having the most significant expression changes during pseudo time trajectory segments **Table 8 (relates to Fig. 5)**. For example, using Moran’s statistical test we compared expression changes from C6 (basal) to C5 (luminal) to identify increased *Psca* and *Mal* gene expression **(Fig. 5A)**. From C6 to C22 (luminal), we identified increased *Upk3a* and *Fmo5* expression **(Fig. 5B)**. Transitioning from C6 (basal) to C11 (squamous cell carcinoma) revealed marked reductions of *Krt5* and *Psca* but increases in *Col12a*, *Thbs2* and *Enpp2* **(Fig. 5C)**. Transitioning from C6 (basal) to C8/0 (fibroblast) revealed increased BGN-*Bgn* expression **(Fig. 5D)**. Finally, transitioning from C6 (basal) to C2 (mesenchymal stem cell) showed heightened *Clec3b*, *Serping1*, *Col1a2* and *Ptx3* gene expression **(Fig. 5E). Table 8, 9, 10** (relates to Fig. 5).

**Fig. 5.**
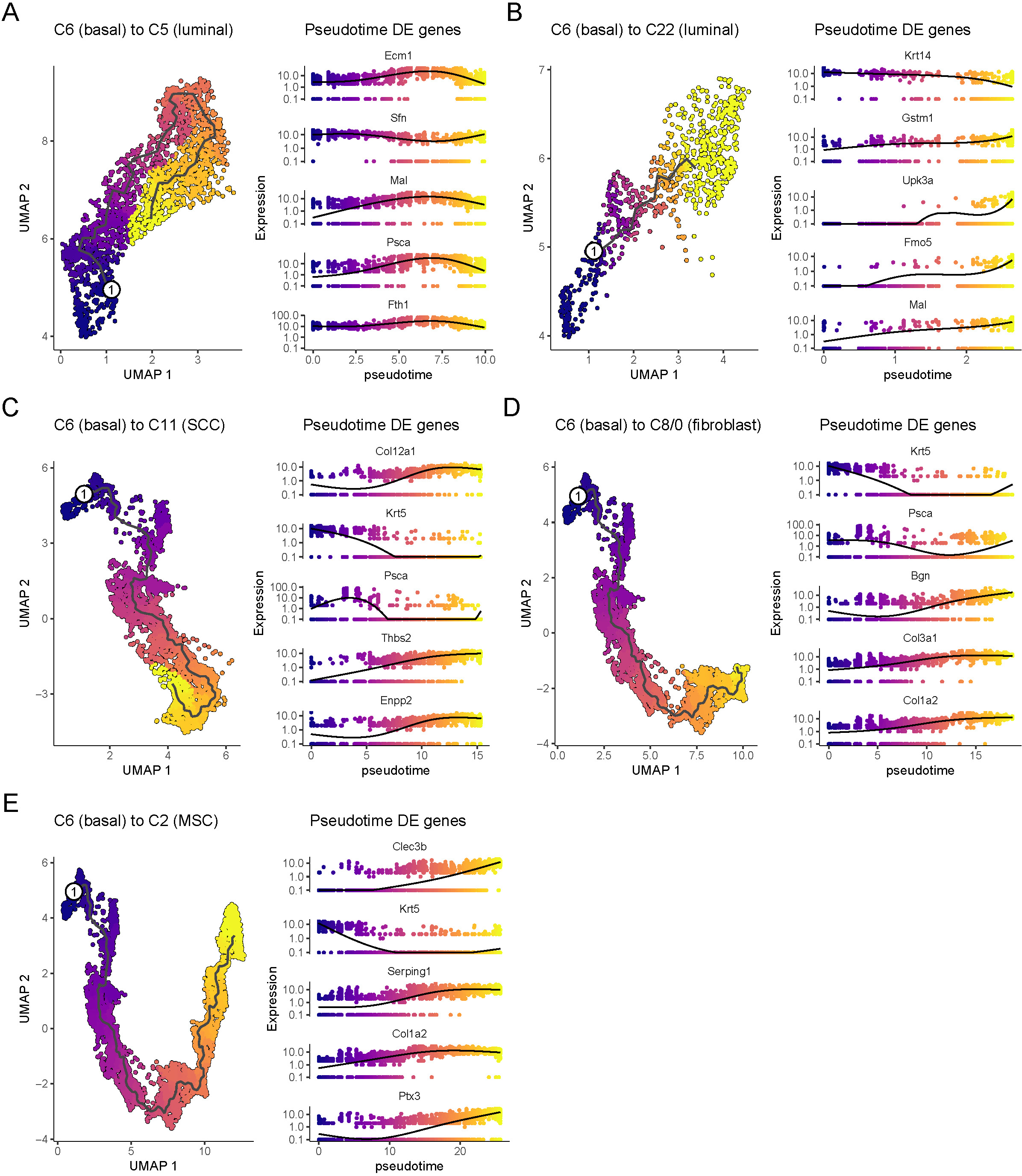
Differentially Expressed Genes (DEGs) during pseudotime. Starting from position 1 (cluster 6, basal), DEGs are shown during pseudotime analysis terminating in (A-B) luminal (cluster 5 and 22), (C) SCC (cluster 11), (D) fibroblast (cluster 0/8), and (E) MSC (cluster 2) compartments. Relates to **Tables 8, 9 and 10**.

*Together, these data show the temporal changes in signature gene expression between tumor cell identities and simulate possible ways transcriptional programs can dynamically change in cell populations to give rise to other lineages. Comparing gene pathways between tumor identities reveals differential pathway enrichments and potential drug sensitivities*.

### Reactome and drug target pathway enrichment in tumor clusters

The tumor clusters identified in this study contain complex and heterogeneous basal, luminal, and EMT marker distribution. This is exemplified by the multi-cluster expression of *Trp63*, *Krt5* (basal), *Krt8* (luminal), and *Timp1* (EMT) throughout the 14 tumor of which several have high Mki67 (Ki67) expression, indicating active cell proliferation **(Fig. 6A)**.

**Fig 6.**
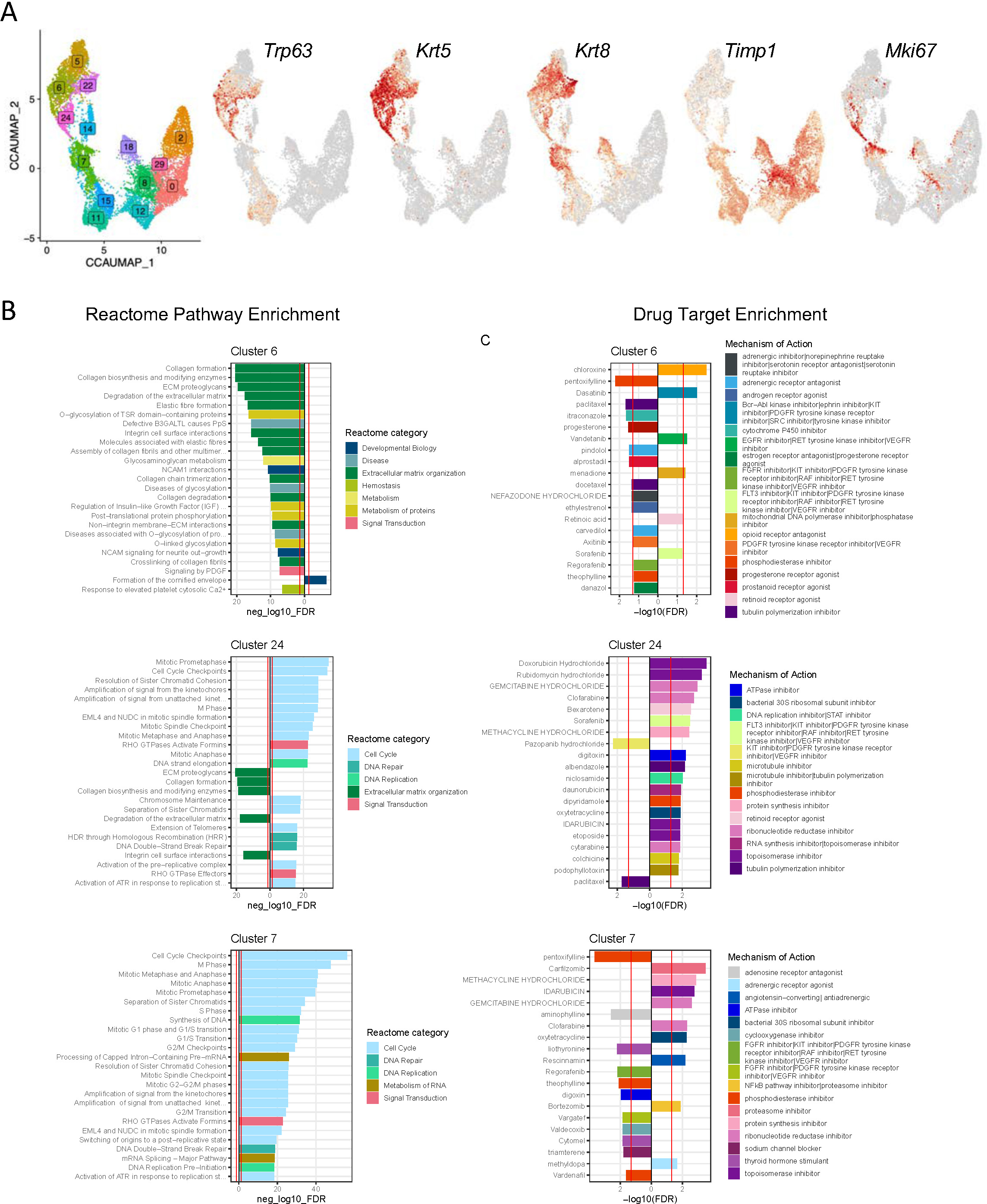
Reactome pathway and drug target enrichment in tumor cell clusters. (A) UMAP plots showing cell clusters and the presence of basal (*Trp63*, *Krt5*), luminal (*Krt8*), EMT (*Timp1*), and cell proliferation (*Mki67*). (B) Reactome pathway enrichment in unique tumor cell clusters (C6, basal urothelial cells; C24, basal urothelial cells; C7, squamous cell carcinoma). (C) Drug target enrichment is shown by clusters (C6, C24, C7). X-axes show multiple testing-corrected significance (-log10(FDR)) of positive and negative enrichment, with vertical red lines marking a significance cutoff of -log10(0.05)**. Relates to Tables 11, 12, and 13.**

Using the information from the trajectory analysis, we generated enrichment plots comparing three tumor clusters with differential lineage marker composition. We assessed the activity of pathways that function through different mechanisms for cluster 6_basal cell of the urothelium, cluster 24_basal cell of the urothelium, and cluster 7_adenocarcioma **(Fig. 6B)**. In each trajectory, multiple hits were identified (x-axis, red lines, p = 0.05) for each trajectory color-coded by the Reactome category as shown in **Table 12** (relates to Fig. 6B).

We observed different gene expression mechanisms in each cell state, with cluster 6 showing dependence upon extracellular matrix organization and PDGF signaling. Cluster 24 showed high expression of cell cycle pathways and activation of RHO GTPase signaling (middle panel), while cluster 7 showed heightened RNA metabolism (lower panel). We then analyzed drug target enrichments for clusters 6, 24, and 7 to determine whether clusters with different transcriptional landscapes may have differing therapeutic sensitivities **(Fig. 6C)**. We cross-matched drug MOAs during progression from node 1 to clusters 5, 22, 11, 8/0, and 2 to generate adjusted p values (column E) related to gene pathway alterations (column G) and drugs that relate to these pathways (column A). **Table 13** (relates to Fig. 6C).

For cluster 6, we identified inhibitors operating by different mechanisms, including chloroxine (opioid receptor antagonist), pentoxifylline (phosphodiesterase inhibitor), and dasatinib (multi-tyrosine kinase inhibitor). Cluster 24 analysis showed enrichment for multiple drugs, including chemotherapeutic (Doxorubicin, Rubidomycin, Gemcitabine), ribonucleotide inhibitors (Clofarabine), retinoid receptor agonists (bexarotene) and kinase inhibitors (Sorafenib). Cluster 7 unique drug enrichments included proteasome inhibitor (carfilzomib), thyroid hormone stimulant (liothyronine), topoisomerase inhibitor (Idarubicin), and sodium channel blocker (Clofarabine).

*These data show that BBN tumor cell clusters have unique molecular landscapes and pathway enrichments, which may dictate distinct drug sensitivities*.

## DISCUSSION

This study investigated tumor heterogeneity in a carcinogen-initiated mouse model of muscle-invasive bladder cancer. Using BBN-induced tumors and derived single-cell RNA-seq data sets, our study has conducted the following investigations: (1) The *Tabula Sapiens* transcriptional atlas and Unicell deep learning were used to establish a detailed molecular landscape of primary BBN-induced tumors. (2) Unicell defined unique tumor signatures, which we have characterized as urothelial, adenocarcinoma, squamous cell carcinoma, fibroblastic, mesenchymal, and epicardial. Within these Unicell-defined signatures, we studied subpopulations with unique combinations of lineage cell markers, including those with known clinical biomarker function [3]. (3) Pseudotime trajectory analysis demonstrated the potential for tumor cell plasticity when initiated from basal cells with known tumor-initiating function, and (4) We applied Reactome pathway analysis to identify transcriptionally defined cell clusters to predict sensitivity to targeted therapies.

Our previous investigation demonstrated the presence of bladder tumor cells with K5-K8 hybrid in lineage marker expression. It used cell-based and *in vivo* transplant assays to show that cell populations enriched with basal cell signatures can progress to form multilineage tumors [11]. Our present study extends these observations by using deep learning to define a detailed molecular landscape and provide evidence that discreet tumor subpopulations are heterogeneous and dynamic.

The data and modeling from our study provide a baseline to assess the functional association of lineage-defined tumor subpopulations with clinical parameters, including tumor progression, response to therapy, and survival. Specifically, we highlight that untreated primary bladder tumors contain a complex array of epithelial, stromal, and hybrid populations that cannot be assigned as basal or luminal in signature. With a panel of basal, luminal, and EMT markers, we identified the presence of 5 strictly epithelial clusters (clusters 5, 6, 14, 22, 24) consisting of basal-luminal hybrid populations and two epithelial populations consisting of a luminal-EMT hybrid signature based on Krt8-Vim co-expression (clusters 7, 18). The remaining transcriptional clusters include nine subpopulations with mesenchymal qualities (clusters 0, 2, 7, 8, 11, 12, 15, 18, 29), including those with heightened expression of transcription factors known to program EMT (e.g., cluster 29). We further identified cluster-specific expression of Chd2 (N-Cadherin) (clusters 8, 18), a marker for neuroendocrine tumor cells and a stimulator of FGFR1. The broad expression of Beta-catenin (Ctnnb1) suggests the possibility of an essential regulatory role in promoting disease progression. Future investigations should assess the impact of acute and prolonged drug treatment on the tumor composition of subpopulations as well as those that may be associated with treatment resistance.

Current clinical practices typically use IHC-based assays for prognostic biomarker detection in solid tumors [3]. Our study used signatures composed of multiple gene biomarker markers for each lineage type, including basal, luminal, and EMT signatures. With a panel of markers, we have provided greater confidence in defining the lineage composition of tumor clusters than using a single marker. The ability to conduct sensitive changes in tumor subpopulations in clinical specimens during progression or treatment will require refinement of IHC grade staining procedures or advances in transcriptional-based assays such as spatial single cell sequencing.

Lineage plasticity is associated with progression and response to therapy in several solid tumors, including prostate cancer, in which tumor epithelia high in AR expression can undergo treatment-induced transdifferentiation to neuroendocrine-like tumor cells in response to therapy [23]. Considerably less is understood about whether progression or treatment-induced changes can occur in the molecular landscape of bladder tumors.

However, select studies have investigated the impact of progression-induced cell-intrinsic changes. For example, in phylogenetic analysis, urothelial and squamous bladder cancer cells have been shown to originate from a common precursor with loss of the transcription factors FOXA1, GATA3, and PPARG being responsible for the maintenance of the urothelial lineage identity [23]. Also, experiments using human bladder cancer cell lines have demonstrated bidirectional switching between epithelial and mesenchymal states [24]. Finally, using a genetic mode, studies have shown that forced expression of MYC, AKT and RB-loss are sufficient to induce the formation of small cell carcinoma (SCCB) from normal urothelial [3]. Such studies are compatible with our trajectory analysis, indicating that lineage identities are not static but dynamic and regulated by cell-intrinsic signaling, progression, and potential therapy. Future studies may focus on (1) the efficiency of each subpopulation to undergo plasticity and lineage change and (2) circumstances in which plasticity is reversible, particularly in response to therapy.

Our study has several limitations. *First*, while we applied a panel of lineage-specific markers, including those with clinical biomarker function, additional regulatory factors may be required to define functional basal-luminal-EMT signatures during progression and in response to therapies. *Second*, BBN tumors progress through many pathological states. However, our data set focuses only on MIBC, and thus, the identified transcriptional landscapes defined here may not represent signatures found in earlier disease states, including NMIBC. *Third*, while the BBN model shows striking transcriptional similarities with human BLCA progression, including significant clinical variants and lineage hybrid tumors, it is unclear whether the signatures identified in this study are present in treating naïve human tumors.

Together, our data justify and rationalize future investigations concerning the changes in tumor sub-populations during progression and in response to therapeutics, including chemotherapy, immune therapy, and other targeted therapies. More advanced analysis may include subpopulation changes during acquired resistance and progression to metastasis.

## LIST OF TABLES

Table 1. Cell cluster numbers (Relates to Fig. 1)

Table 2. Cell type numbers (Relates to Fig. 1)

Table 3. Unicell classification genes (Relates to Fig. 2)

Table 4. Unicell cell type DEGs (Relates to Fig. 2)

Table 5. Unicell cell type DEG Pathway Enrichment (Relates to Fig. 2)

Table 6. Unicell cell type DEG Drug target Enrichment (Relates to Fig. 2)

Table 7. P-values for genes of interest along pseudotime trajectory branches relates to Fig. 4

Table 8. Pseudotime trajectory DEGs (Relates to Fig. 5)

Table 9. Pseudotime trajectory DEG Pathway Enrichment (Relates to Fig. 5)

Table 10. Pseudotime trajectory DEG Drug Target Enrichment (Relates to Fig. 5)

Table 11. Cluster DEGs (Relates to Fig. 6)

Table 12. Cluster DEG Pathway Enrichment (Relates to Fig. 6)

Table 13. Cluster DEG Drug Target Enrichment (Relates to Fig. 6)

## SUPPLEMENTAL MATERIALS

**Fig. S1.** UMAP plots showing coexpression of Krt14 and Krt8 positive cells. UMAP plots and heat maps showing Krt14 (red) and Krt8 (green) expressing cells. Yellow depicts Coexpression (Relates to Fig. 3D).

**Fig. S2.** The expression of basal genes is shown as a (A) UMAP plot and (B) individual genes (Relates to Fig. 3A).

**Fig. S3.** The expression of luminal genes is shown as a (A) UMAP plot and (B) individual genes (Relates to Fig. 3B).

**Fig. S4.** The expression of mesenchymal genes is shown as a (A) UMAP plot and (B) individual genes (Relates to Fig. 3C).

