## Supplemental Figures for "Mapping of Multilineage Tumor Cell Populations in Mouse Bladder Cancer"

Fig. S1

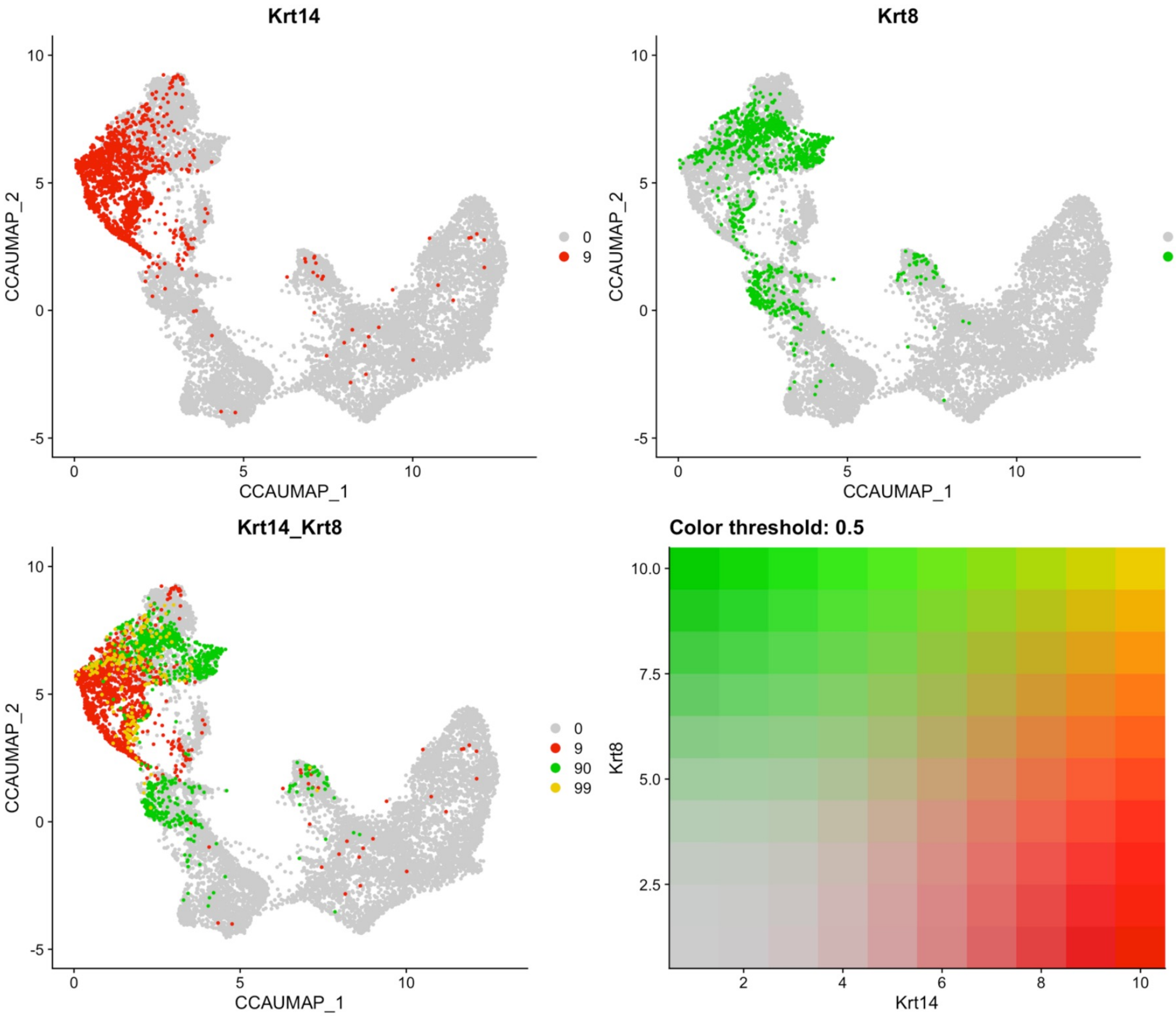

Krt14
Krt8
Krt14_Krt8
Color threshold: 0.5
CCAUMAP_1
CCAUMAP_2
0
9
90
99
Krt8
Krt14

Fig. S2

A **All basal gene co-expression**

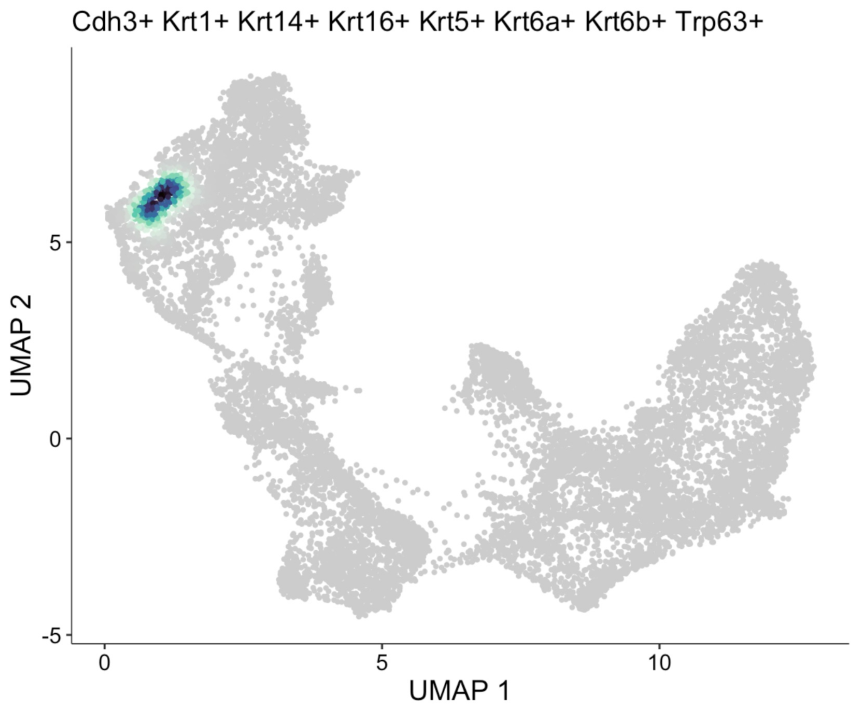


B **Individual basal genes**

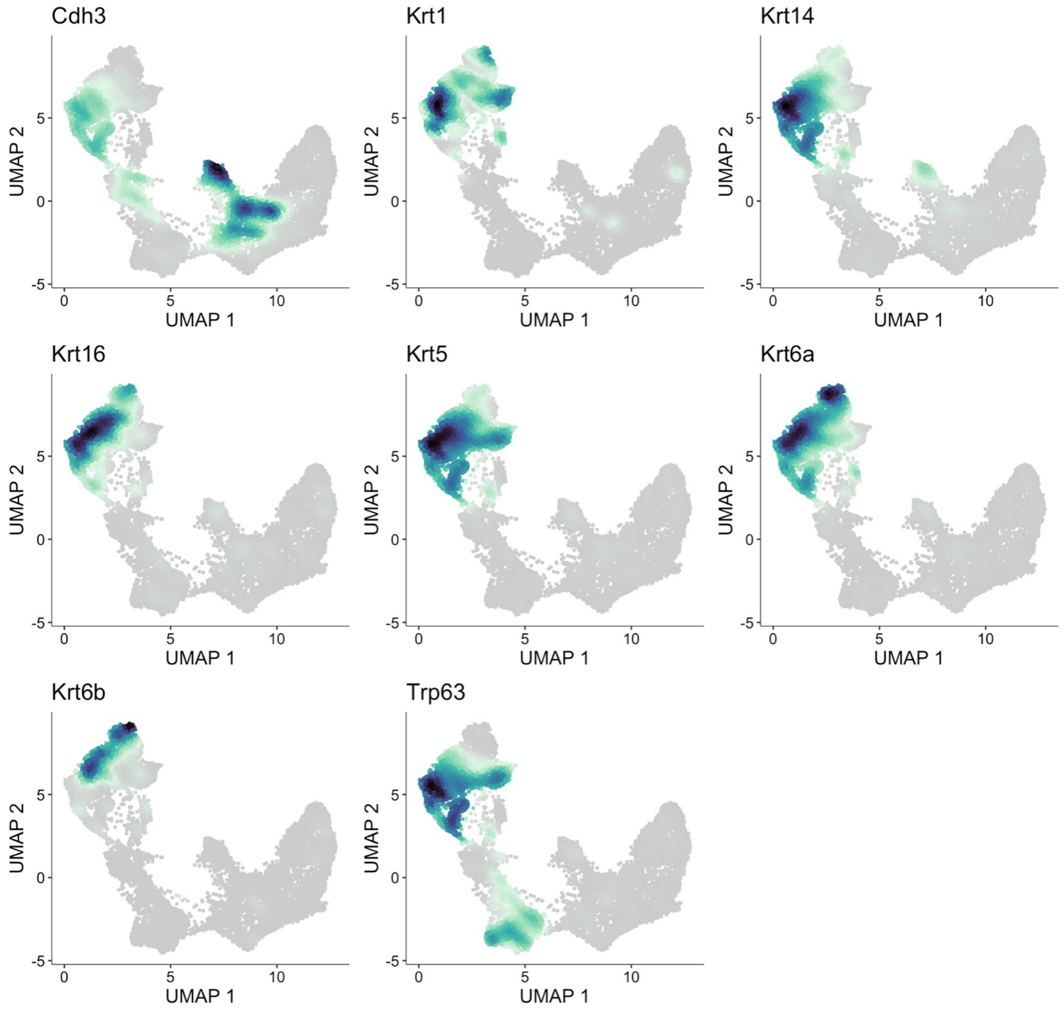

Fig. S3

A

**All luminal gene co-expression**

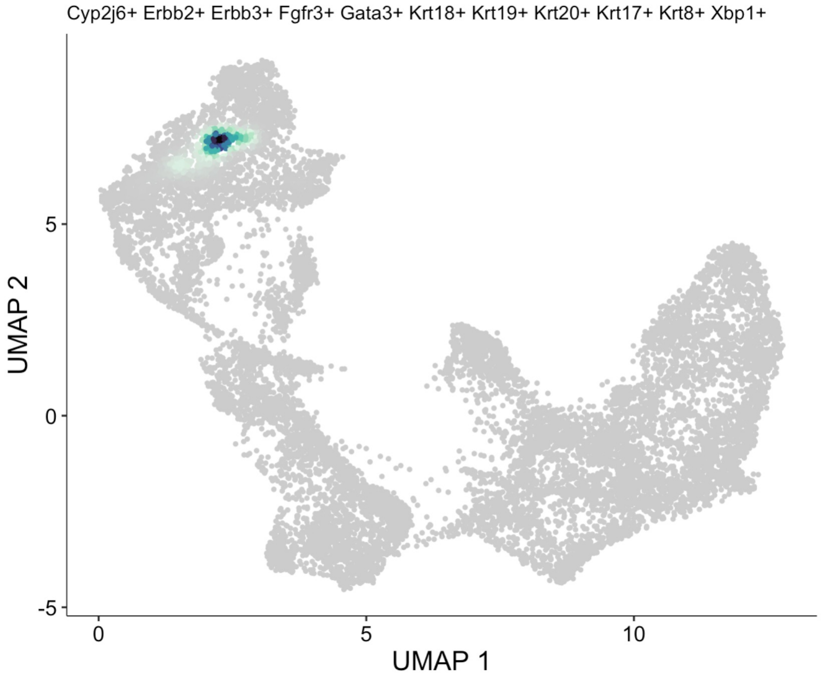


B

**Individual luminal genes**

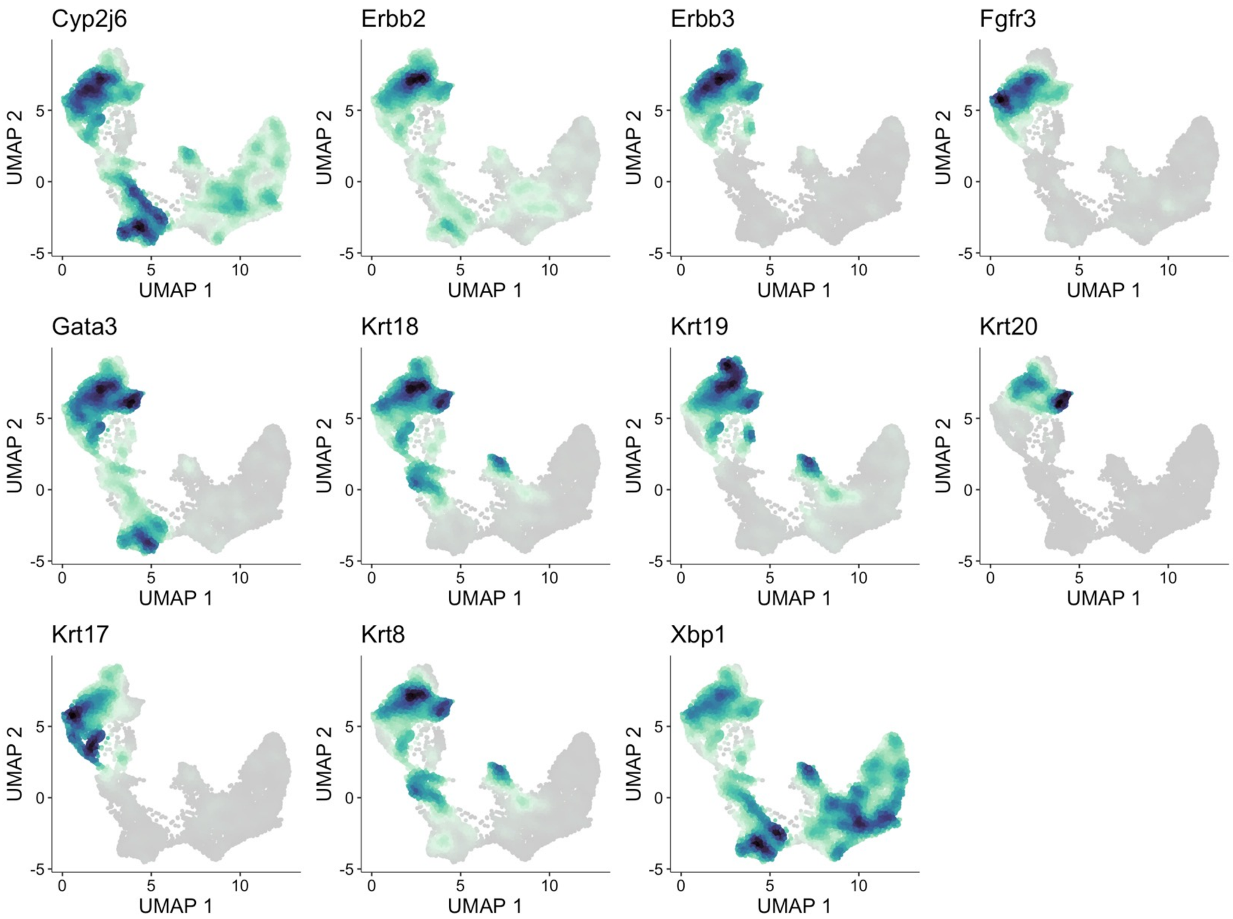

Fig. S4

A **All EMT gene co-expression**

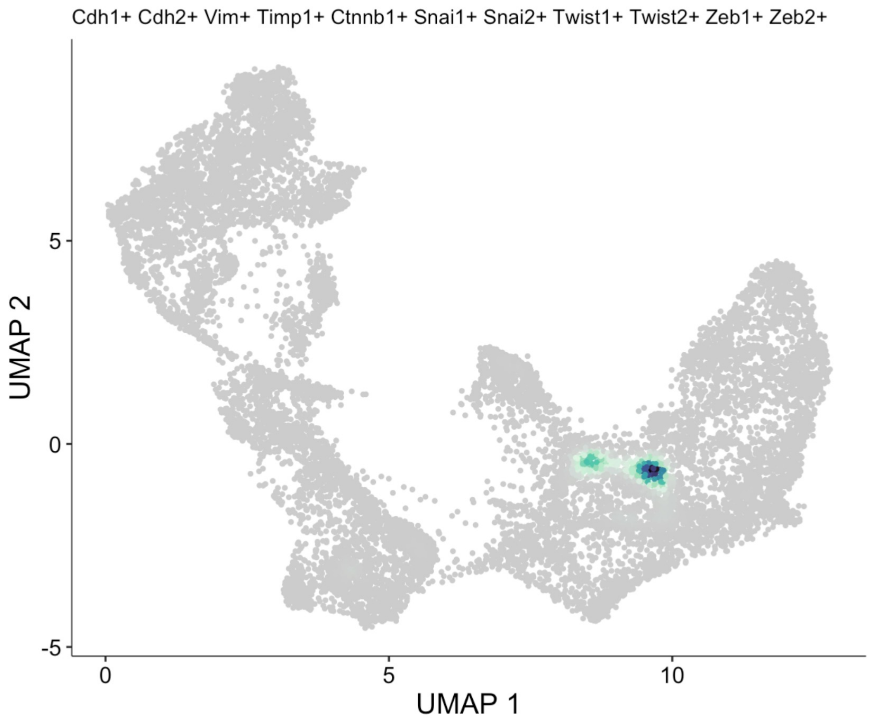


B **Individual EMT genes**

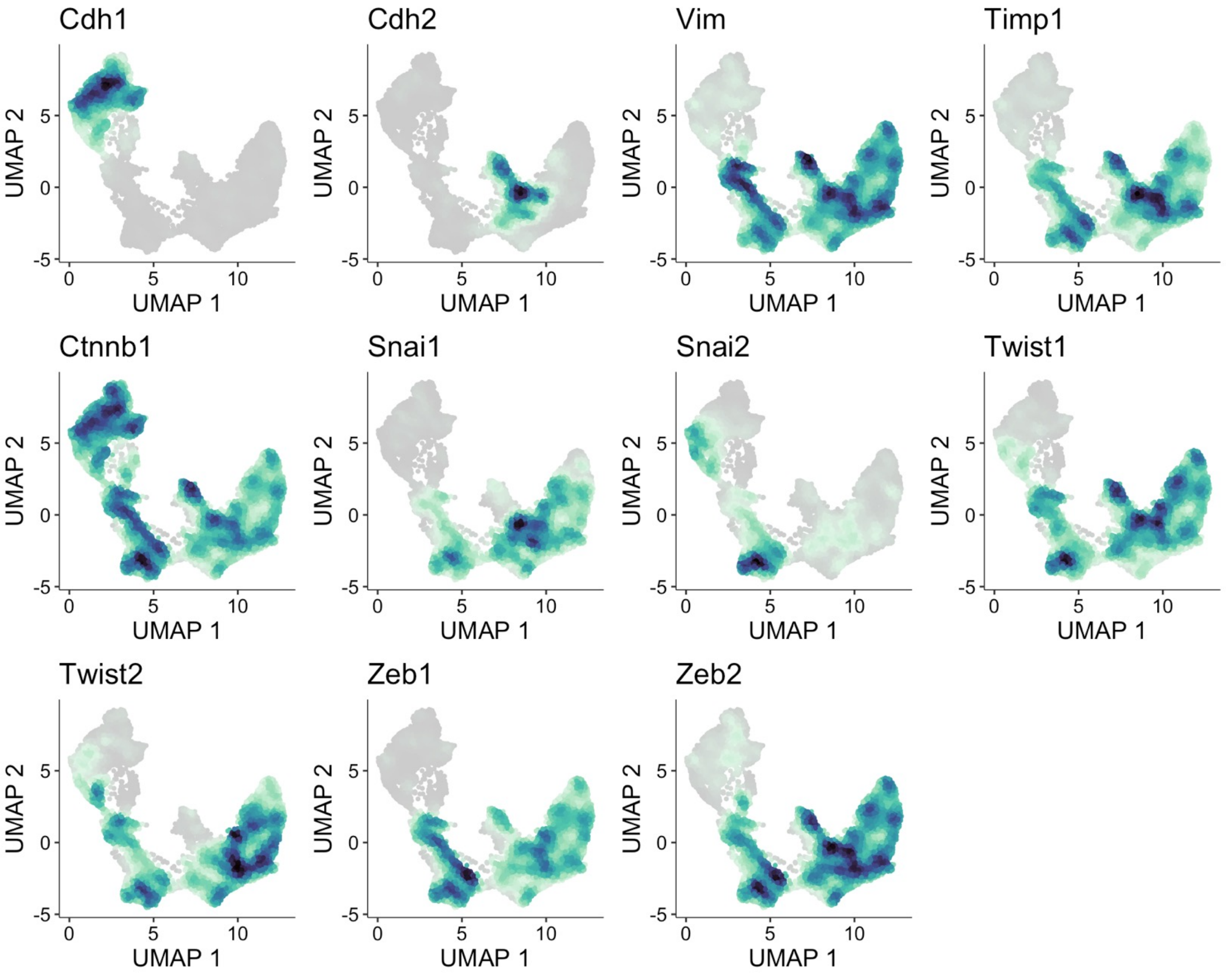
